# TIPP-SD: A New Method for Species Detection in Microbiomes

**DOI:** 10.1101/2025.08.27.672749

**Authors:** Chengze Shen, Eleanor Wedell, Mihai Pop, Tandy Warnow

## Abstract

In this study, we present TIPP-SD (i.e., TIPP for Species Detection), a new technique for species detection in a microbiome sample. TIPP-SD uses a modified version of TIPP3, which is a recently developed abundance profiling tool based on maximum likelihood phylogenetic placement into marker gene taxonomies. TIPP-SD depends on a parameter (i.e., “threshold”) for the required support for species detection, thus allowing us to compute a precision-recall curve as we vary this parameter. In comparing the precision-recall curves for TIPP-SD, TIPP3, Kraken2, Bracken, and Metapresence, we find that TIPP-SD improves on the other methods with respect to accuracy under conditions where some species occur in low abundance, or where there is sequencing error. Under easier conditions, TIPP-SD is close to the best of these methods. Finally, although TIPP-SD is slower than Kraken2 and Bracken, it is still fast enough to be used on large datasets.

## 1 Introduction

Abundance profiling and species detection are two related problems in microbiome analysis. Species detection aims to list the species found in an environmental sample, while abundance profiling aims to estimate the distribution of species (or genera, families, etc.) within the sample. Each of these problems has broad applications and together they provide insight into questions related to ecology, evolution, and human health [4, 29, 31, 19, 21].

Some of the methods for these problems include Bracken [23], Kraken2 [42], krepp [34], MEGAN4 [12], Metabuli [16], MetaPhlan4 [3], Metaphyler [20], mOTU [32], sylph [36], the TIPP family of methods [27, 35, 38], and YACHT [17]. While methods for abundance profiling can be used for species detection by setting a threshold above which a species is considered to be detected, the selection of the appropriate threshold is itself a non-trivial question. In addition, because the goal of abundance profiling is an estimate of the distribution (relative frequencies) at a given taxonomic level, such as species, abundance profiling methods can afford to ignore very low abundance taxa, as setting their frequency to zero is unlikely to hurt the overall accuracy by more than a very small amount. However, such a strategy can reduce their utility for species detection. Other issues in using abundance profiling for species detection are discussed in [33], who specifically emphasize the importance of detecting species that are present in very low frequencies (i.e., “rare taxa”), and argue that abundance profiling methods are insufficient for this purpose. For multiple reasons, therefore, methods specifically designed for species detection are of interest.

In this study, we propose TIPP-SD, a new technique for species detection. While TIPP-SD builds on the basic algorithmic design in TIPP3 [38], in order to achieve high precision and recall for species detection, it modifies the TIPP3 technique in several ways.

We compare TIPP-SD to the naive use of TIPP, and show that TIPP-SD provides superior accuracy for species detection. We also compare TIPP-SD to Bracken [23], Kraken2 [42], and Metapresence [33]. We find that TIPP-SD has very good accuracy, with improved accuracy in many conditions. We also find that TIPP-SD is slower than Kraken2 and Bracken, but is still fast enough to be used on very large datasets.

The rest of the paper is organized as follows. We provide background information in Section 2, including details about TIPP3. We present TIPP-SD in Section 3. We present the experimental design in Section 4, and the results are provided in Section 5. A discussion of the trends observed is provided in Section 6. The supplementary materials provide additional results and details.

## 2 Background

We provide some background here, including a description of the TIPP3 method for abundance profiling.

### 2.1 Abundance profiling methods based on marker genes

The goal of abundance profiling is to obtain a good estimate of the distribution of the species, genera, families, etc., in a given microbiome sample. While this problem can be considered at different taxonomic levels, for the purpose of this discussion, we will assume the interest is in estimating the abundance at the species level.

The input is typically a set of reads, though in some cases contigs may also be used, which could be based on amplicon sequencing (e.g., just one gene) or metagenomics (from across the genome). Because the goal is to produce a good estimate of the relative frequencies of species, abundance profiling methods must consider factors that distort the estimates of relative abundance: the genome length and whether genes appear in multiple locations within a genome [26, 22]. Because of these challenges, *some* methods for abundance profiling are based on filtering the input reads to those that are derived from *marker genes*, which are genes that are expected to be single-copy and universal. Examples of such methods, which are referred to as “marker gene-based methods”, include Metaphyler [20], MetaPhlan [39], mOTUs [32], and the TIPP family of methods [27, 35, 38]. Of these, TIPP3 [38] has been shown to have the best accuracy for abundance profiling, especially when working with reads generated by sequencing technologies that have high error rates (indels or substitutions), such as Nanopore [40] and PacBio [30]. As shown in [38], restricting Kraken2, Bracken, and Metabuli to the reads that are mapped to the TIPP3 marker genes *improved* the abundance profiling distributions that each method produced. Furthermore, even after this modification to Kraken2, Bracken, and Metabuli, TIPP3 was still more accurate than Kraken2, Bracken, and Metabuli in nearly all conditions; the exceptions were limited generally to Illumina reads (which have very low error rates). TIPP3 was also more accurate than Metaphlan4 [3], which is another marker-gene based method.

Because TIPP3 had high accuracy for abundance profiling, we seek in this study to build on its technical approach, but modify it so as to obtain high accuracy for the species detection problem.

### 2.2 TIPP3

We begin with a description of the high-level approach of the TIPP3 method; see [38] for full details. The (publicly available) TIPP3 reference package includes multiple sequence alignments and taxonomies for each marker gene in the package. TIPP3 uses BLASTN [1] to map reads to the TIPP3 marker gene reference package. Any read that does not map to a marker gene is discarded. Each read that maps to a marker gene is added into the corresponding marker gene alignment using WITCH [37].

The next step in the analysis places each read into the taxonomy for that marker gene, using a phylogenetic placement method based on maximum likelihood, such as pplacer [24], EPA-ng [2], or BSCAMPP [41] (a divide-and-conquer approach that enables pplacer and EPA-ng to be used on large trees). Of these phylogenetic placement methods, pplacer is the most accurate but cannot be used on large trees very easily. EPA-ng is faster than pplacer and slightly less accurate. By default, TIPP3 uses pplacer-taxtastic [5, 9], a way of using pplacer that allows it to run on moderately large trees.

The output from the phylogenetic placement step is a list of the top edges in the tree (using maximum likelihood as the criterion), along with the relative support for each edge. TIPP3 picks the edge in the rooted taxonomy so that the support for the read belonging to the clade below that edge is at least *B*, where *B* is an input parameter set to 0.95 or 0.9 depending on whether EPA-ng or pplacer is used for phylogenetic placement. The selected edge determines the taxonomic labels for the read: it may be given a species label, or it may only be placed at a genus or higher level. Finally, the taxonomic information across all the reads that map to marker genes is aggregated to form an abundance profile at each taxonomic level.

A fast version of TIPP3 (TIPP3-fast) makes two adjustments to the TIPP3 design in order to reduce the runtime, with a small increase in error: instead of using WITCH to add reads into the marker gene alignment it uses BLASTN, and instead of using pplacer-taxtastic for phylogenetic placement it uses BSCAMPP with EPA-ng.

## 3 TIPP-SD

We now describe TIPP-SD. We describe it here for species detection, but note that the algorithm can be generalized to detect other taxonomic levels as well.

Recall that TIPP3 has a publicly available reference package of reference taxonomies and multiple sequence alignments, one for each of its marker genes; TIPP-SD uses these in its analyses. Recall also that TIPP3 places reads into these taxonomies using a maximum likelihood phylogenetic placement method, and these placements return statistical support values on the edges. TIPP3 considers the statistical support returned by these methods, and only places a read into an edge if the clade below it has support at least *B* (where *B* is an input parameter, set by default to be either 0.9 or 0.95, depending on which phylogenetic placement method is used).

The first difference between TIPP-SD and TIPP3 is that we do not use *B* to inform the placement of each query sequence. Second, we have a fast but accurate way of performing the phylogenetic placement. Recall that TIPP3-fast (the fast version of TIPP3) uses BSCAMPP with EPA-ng to speed up the read placement step. BSCAMPP using pplacer as the base method is more accurate than BSCAMPP with EPA-ng, with only a small increase in running time [41]. In this study, we explore using BSCAMPP with pplacer (i.e., BSCAMPP(p)) for read placement and BLASTN for read alignment. This modified technique aims to improve accuracy in phylogenetic placement of reads into the taxonomies and sacrifies a little accuracy for the read alignment, which adds the reads into the multiple sequence alignment for the marker gene, without increasing the runtime too much.

The next change between TIPP-SD and TIPP3 is how we use BSCAMPP(p), the maximum likelihood phylogenetic placement method, to classify a read at a species level, assuming sufficient statistical support. Given a single read that is mapped to the marker gene *g*, BSCAMPP(p) returns support values for the top seven (by default) edge placements in the tree, based on their likelihood values. If none of these are at the species level, then the read will not be classified at the species level. Otherwise, we use the support value returned by BSCAMPP(p) for the placement of the read into the taxonomy at the species level as the “confidence score” for this species assignment, and use this in the next step.

Given this information, we consider two techniques for using TIPP-SD for identifying the set of species available in the sample: *marker vote* and *marker confidence*, each of which depends on a user-specified threshold *T*, as we now describe.

### Marker vote

Each read that is assigned a species label using the technique from the previous paragraph is also mapped to a marker gene, and we consider the marker gene to have therefore voted for that species. Thus, after aggregating results from all mapped reads, we will have a collection of species, each with a list of marker genes that voted for the species. We let *vote*(*s*) denote the number of votes for the species *s*.

In the TIPP3 reference package, a species may not appear in all the marker gene trees, depending on the quality and completeness of the genomes for the species. Therefore, for each species *s*, we note the number *m*_*s*_ of marker genes in which *s* appears. The ratio 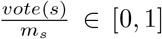 then represents the fraction of marker genes that vote for the presence of species *s*. By setting a threshold *T* and checking if 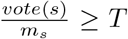, we can determine if a species *s* is detected. Thus, the output of marker vote depends on the threshold *T*.

### Marker confidence

Here we describe a technique for assigning a confidence for a species being present that takes into account the statistical support values for a read to belong to each of the species, as provided by BSCAMPP(p) during the read placement stage. Thus, we do not select the single best species assignment for a read, but consider all the assignments with positive support, and then normalize appropriately. We describe this technique in detail below.

For a given gene *g* and species *s* contained in the gene *g*, we let *c*(*g, s*) be the sum of the confidence scores for *s* across all the reads mapped to gene *g* that have non-zero support for *s*, divided by the number of such reads. We sum *c*(*g, s*) across all the *m*_*s*_ genes *g* that contain *s*, and divide this by *m*_*s*_ to obtain the final confidence score *C*_*s*_ for species *s*. By setting a threshold *T* for confidence, we will say that *s* is detected if *C*_*s*_ ≥ *T*. Thus, the marker confidence output depends on the threshold *T*.

## 4 Experiment Design

In our experiments, we explore different methods for identifying species on datasets generated with varying types of sequencing technologies, where we know the true list of species present in the sample. This allows us to quantify both precision and recall for each analysis, and to explore the conditions under which each method is accurate.

### 4.1 Methods

We compared TIPP-SD and its variants to TIPP3, Kraken2 [42], Bracken [23], and Metapresence [33]. Here, we briefly describe each method and the databases they use in comparison to TIPP-SD.

#### Kraken2 and Bracken

Kraken2 and Bracken are both taxonomic identification tools that assign taxonomic labels to input reads. They can also be used for abundance profiling, and we used them for species detection based on their reported number of reads assigned to each species. Kraken2 uses its database and does k-mer searches for read classification. We used its published database *maxikraken2-PlusPF* on June 5th, 2023, with 24,320 unique species. This number of species is close to the number of species (25,509) used in the TIPP3 reference package. Bracken uses the Kraken2 output and modifies the reads for its report, hence using the same database.

#### Metapresence

Metapresence is specified for species detection by mapping metagenomic reads to genomes. It pre-builds a Bowtie2 database [18] on a fixed set of genomes, and uses this database for species detection in input reads. In the Metapresence study [33], the authors did not provide a pre-built database. Hence, for this study, we built a customized Bowtie2 database using the procedure described in Metapresence. We made sure the comparison is fair between TIPP-SD and Metapresence by restricting the TIPP3 reference package to the same set of species in the Bowtie2 database. More details can be found in Section 5 regarding the creation of the customized Bowtie2 database.

### 4.2 Benchmark datasets

A summary of the datasets we used and their properties can be found in Table 1. We provide more details for the read simulation and each dataset in the following paragraphs.

**Table 1:** Properties of simulated datasets.

| Dataset | Type | Number of species | Number of reads | Mean length | Avg coverage |
| --- | --- | --- | --- | --- | --- |
| 50 Known | Illumina | 50 | 10,500,910 | 150 | 20 |
|  | PacBio | 50 | 1,047,884 | 3006 | 20 |
|  | Nanopore | 50 | 184,327 | 4033 | 5 |
| 1000 Known | Illumina | 1000 | 13,795,579 | 150 | 1 |
|  | PacBio | 1000 | 1,418,197 | 2925 | 1 |
|  | Nanopore | 1000 | 921,633 | 4036 | 1 |
|  | Nanopore | 1000 | 9,216,327 | 4032 | 10 |
| CAMI-II Marine | Illumina | 301 | 33,301,262 | 150 | 2 |
|  | PacBio | 301 | 1,641,591 | 2968 | 2 |

#### Sequencing models

We used read simulators to generate Illumina, PacBio, and Nanopore reads of each given set of genomes, as described below (except for CAMI-II datasets, which are available online). We used `art_illumina` (v2.5.8) [11] for generating Illumina short reads with the HS25 error model (fixed 150bp in read length, ~0.1% substitution rates). We used `PBSIM` [28] for generating PacBio long reads with the CLR sequencing error model, with an average read length of 3000bp and a minimum length of 400bp to understand the impact of sequencing error. The CLR model has an average 78% sequencing accuracy, with 3.23% substitution rate, 10.53% insertion rate, and 3.98% deletion rate (obtained by aligning simulated reads to reference using LAST [14]), We used `Nanosim` (v3.2.2) [43] with a pretrained metagenome model of bacteria community to simulate Nanopore reads with a total error rate of 11.3% (3.9% substitution, 3.2% insertion, and 4.2% deletion rates).

#### 50 known species

We first used the same known species datasets from the TIPP3 study [38], with 50 known species in Illumina, PacBio, and Nanopore reads. “Known” means that genomes of the species (not necessarily the same exact genome) are present in the reference package or database of both TIPP-SD and Kraken2/Bracken. These datasets have a coverage of 20 for all included genomes except for Nanopore reads, which have a coverage of 5. Note that the 50 species have an even distribution of abundance.

#### 1000 known species

We also generated a new dataset with 1000 unique species that are also known to TIPP-SD and Kraken2/Bracken, with Illumina, PacBio, and Nanopore reads. Each of the unique species has genome data present in the reference package or database of both TIPP-SD and Kraken2/Bracken. We set the coverage of read simulations to 1 to imitate low species presence in a microbiome sample. We also generated a separate set of Nanopore reads with a coverage of 10 to study whether higher coverage could improve the methods’ performance in recall and/or precision of species detection. Note that the 1000 species also have an even distribution of abundance; thus, each species is expected to appear in 0.1% of the sample.

#### CAMI-II marine dataset

Additionally, we included the Illumina and PacBio reads from the CAMI-II marine dataset, replicate 1 [25]. These two datasets were also studied in [38]. Replicate 1 contains 301 species with non-zero abundance, which are considered for species detection. This dataset has a wide range of abundance levels, and the ten species with the lowest abundance range from 0.0000004% to 0.007% (in contrast, the top 10 species have abundances ranging from 2.4% to 14.4%).

### 4.3 Evaluation metrics

We evaluated each method by its ability to detect species that should be present in the input reads, with precision and recall. Precision is defined as 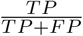, where *TP* is the number of species correctly identified and *FP* is the number of species falsely detected. Recall is defined as 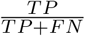, where *FN* is the number of species that should be detected but are not recovered by the method. We also examined F1 score, which is defined as 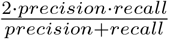 and takes both precision and recall into consideration. Precision, Recall, and F1 score are all in the range of 0, 1].

To obtain the precision-recall curve of a method, we first extracted the method’s taxonomic identification results. We then used a support threshold to determine if a species was present, and varied the support thresholds to obtain precision-recall curves. The precision-recall curves are computed as follows:

- For TIPP-SD used with Marker Vote: the threshold *T* is a value between 0 and 1, and we say the species is detected if the marker vote (a value between 0 and 1) exceeds *T*; the corresponding approach is used with Marker Confidence. By varying *T*, we obtain the precision-recall curve. We also set two default values for *T* in Experiment 1, and indicate these in the figures as red stars.
- For Bracken and Kraken2: we use an integer threshold *X*, and report a species as present if its read count exceeds *X*. By varying *X* between 1 and the largest observed value, we obtain precision-recall curves. Bracken and Kraken2 do not have default thresholds for species detection.
- For Metapresence: this method uses two metrics they define called BER (Breadth-Expected breadth ratio) and FUG (Fraction of Unexpected Gaps) that range from 0 to 1. We consider the pair (*x, y*), where *x* is the threshold for BER and *y* is the threshold for FUG, and retain only the reported species by Metapresence if their BER and FUG scores exceed both thresholds. Then, we can compute the corresponding precision and recall value for the pair (*x, y*). We repeat this process for all pairs of (*x, y*), *x* ∈ 0, 1], *y* ∈ 0, 1], with a step of 0.01, and obtain a collection of (precision, recall) pairs. We sort these values by recall in ascending order and then by precision in descending order, and plot them as the Metapresence precision-recall curve.

## 5 Results

### 5.1 Experiment overview

In Experiment 1, we explored different strategies for species detection with TIPP-SD, as described in Section 3, and selected the best variant and two reporting thresholds for TIPP-SD species detection. In Experiment 2, we explored the impact of filtering input reads [38] to use with Bracken and Kraken2 for species detection, and compared both methods to TIPP-SD with respect to precision, recall, memory, and runtime usage. In Experiment 3, we evaluated Metapresence and TIPP-SD and compared their precision, recall, memory, and runtime usage. Finally, we presented a case study that examines the potential causes of false positives with Kraken2, Bracken, and TIPP-SD, using the 1000 known species datasets.

When comparing each external method to TIPP-SD, we ensured that the reference database/package contains all species we want to detect. In the case of Kraken2 and Bracken (Experiment 2), we selected only species for detection in the intersection of the Kraken2 and TIPP3 databases. In the case of Metapresence (Experiment 3), we only ran it on the three datasets with 1000 known species and built a Bowtie2 database of 2000 genomes. The 2000 genomes contain the genomes of 1000 species we want to detect, and an additional 1000 genomes which have 80-95% average nucleotide identity (ANI) to at least one of the known genomes. ANI was computed using FastANI [13]. We also restricted the TIPP3 reference package to the same genome set when comparing TIPP-SD to Metapresence. The reason why we want this consistency is because of the findings from [6], which point out that database size positively correlates with the loss of resolution at the species level. We also tried creating a Bowtie2 database for Metapresence that would include all unique species from the TIPP3 reference package (~25k unique species), but it timed out after 7 days with 16 cores and 512 GB of memory.

### 5.2 Experiment 1: Species detection strategies with TIPP-SD

#### 5.2.1 Marker vote vs. marker confidence

Figure 1 compares TIPP-SD using *marker vote* or *marker confidence* as the strategy to detect species from input reads.

**Figure 1.**
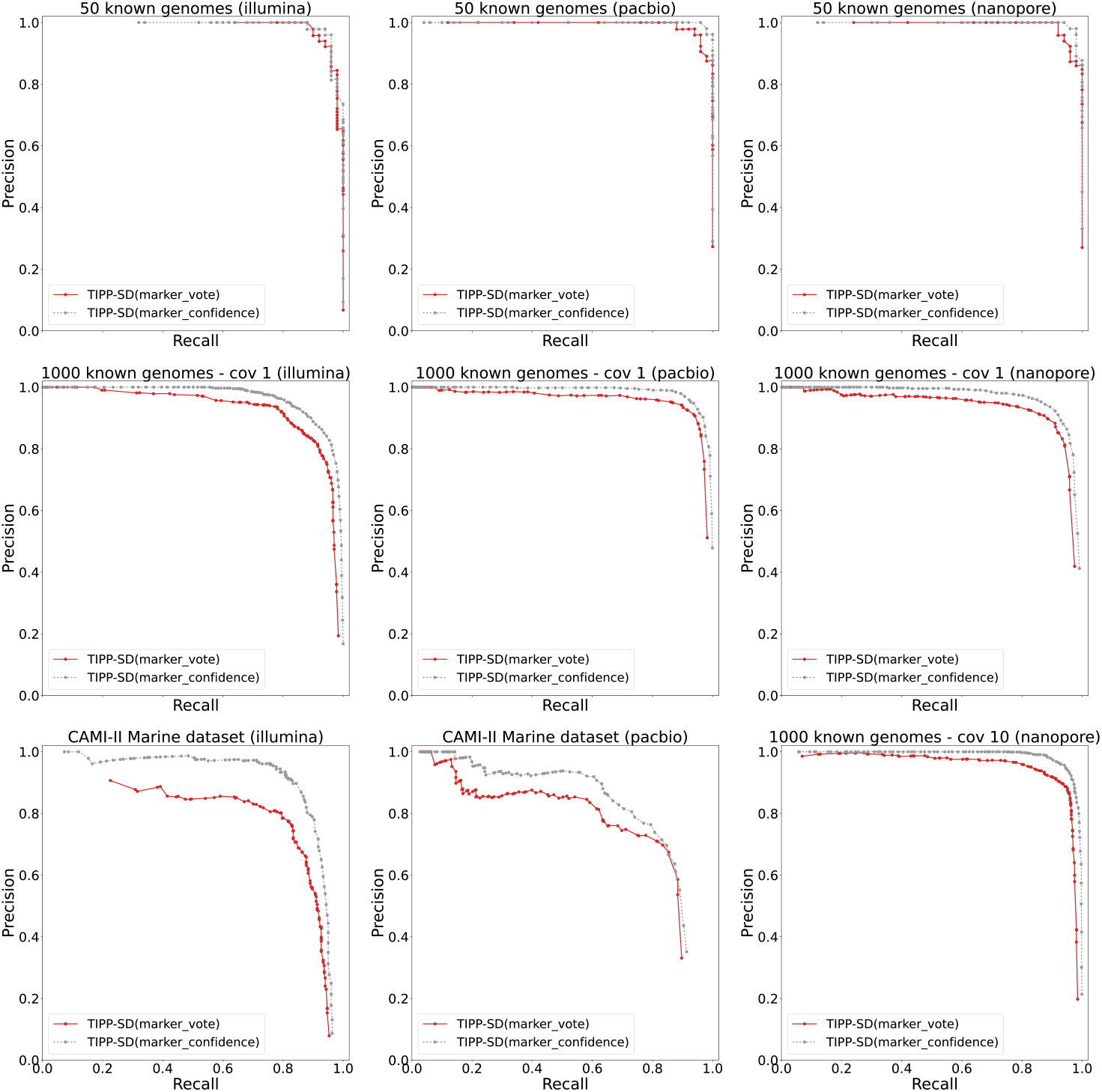
Precision-recall curves of TIPP-SD using either marker vote or marker confidence on nine different benchmark datasets. TIPP-SD is given reads generated from each of the datasets under one of three sequencing technologies (Illumina, PacBio, or Nanopore), as indicated in the name of the dataset above the subfigure. All reads provided to TIPP-SD are from genomes belonging to species also present in TIPP-SD’s reference package. The datasets whose name includes “cov 1” have coverage 1 and those whose name includes “cov 10” have coverage 10. This figure shows that TIPP-SD based on marker confidence produces higher recall than marker vote, and has better precision at each recall value.

Although the difference in the 50 known species datasets between the two strategies is small, using marker confidence achieved higher recall than marker vote, and for a given recall value it also had higher precision. On the 1000 known species datasets, TIPP-SD with marker confidence had better precision and recall than TIPP-SD with marker vote. The same observations hold for the CAMI-II dataset, on both Illumina and PacBio reads. Based on these results, henceforth we only report TIPP-SD using marker confidence.

#### 5.2.2 TIPP-SD variants

We compared TIPP-SD to two variants by changing the alignment and placement methods for reads, in all cases, using marker confidence for species detection. Recall that TIPP-SD uses BLASTN for read alignment and BSCAMPP(p) for phylogenetic placement of reads into the taxonomy for the marker gene. Variant 1 uses WITCH for aligning reads and pplacer-taxtastic for placement (expected to be more accurate but slower), and Variant 2 uses BLASTN for aligning reads and BSCAMPP with EPA-ng for placement (expected to be faster but less accurate).

As seen in Figure S2, Variant 2 was the least accurate method. The comparison between TIPP-SD and Variant 1 shows that Variant 1 had a small accuracy advantage on the 50-genome conditions but they were tied all other conditions. On the other hand, TIPP-SD was much faster and had lower memory usage than Variant 1 (Figures S3 and S4), typically by 1-2 orders of magnitude. Given that the accuracy improvement obtained by Variant 1 over TIPP-SD was small and the increase in runtime and memory usage was very large, we selected TIPP-SD for species detection. Henceforth, we report results using its settings in subsequent experiments.

#### 5.2.3 Selecting default detection threshold

We examined the F1 score with respect to marker confidence for all datasets and showed the comparisons in the supplementary materials (Figure S5). Based on the F1-marker confidence curve, we decided that a “conservative” threshold of value ~0.2 achieves a good F1 score over most conditions for TIPP-SD. For higher sensitivity (recall), TIPP-SD can use a “sensitive” threshold of value ~0.12. We show the precision and recall values of TIPP-SD based on the two thresholds in the following experiments, alongside the precision-recall curve.

### 5.3 Experiment 2: Comparison to TIPP3, Bracken, and Kraken2

This experiment compared TIPP-SD to TIPP3, Bracken, and Kraken2 used for species detection, all of which require the use of a threshold to determine which species are included in the list. Thus, the comparisons were based on precision-recall curves. All methods were compared on the nine different model conditions we explored.

#### 5.3.1 Comparison to TIPP3

We compared TIPP3 to TIPP-SD on all the datasets. Results on the full set of conditions are shown in Supplementary Materials Fig S1, and three conditions are shown in Fig 2. By varying the threshold for abundance, we obtained precision-recall curves for TIPP3. As seen in Fig 2, TIPP3 had poorer accuracy (in particular, lower recall) than TIPP-SD. TIPP3 also has poorer accuracy than TIPP-SD for all 1000 known species datasets, and matches TIPP-SD only on the 50 known species datasets.

**Figure 2.**
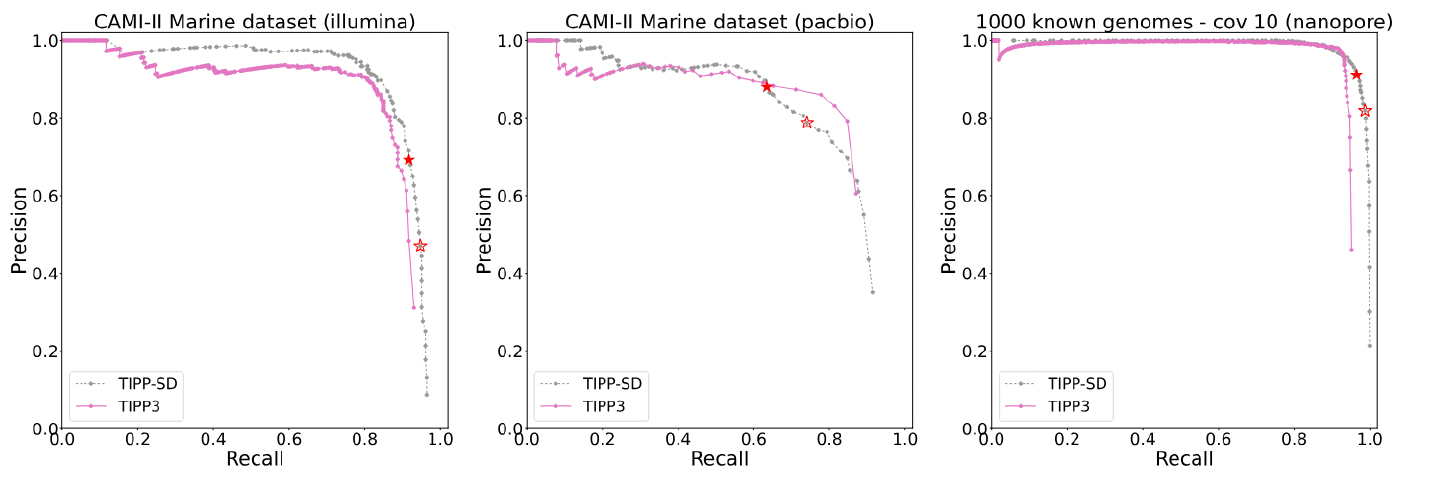
Precision-recall of TIPP-SD compared to TIPP3. TIPP3’s precisionrecall curve is obtained by setting thresholds on the abundance level, and only species with an abundance greater than the threshold are considered as “detected” by TIPP3. TIPP3 is less accurate than TIPP-SD on these datasets, and on the other datasets with at least 1000 species (see Fig S1).

#### 5.3.2 Comparison to Kraken2 and Bracken

We studied the impact of filtering metagenomic reads based on the TIPP3 marker genes as input to Bracken and Kraken2. Bracken and Kraken2 had better abundance profiling accuracy with filtered reads than all reads in the TIPP3 study [38]. However, filtering reads had a mixed impact for species detection with the two methods (Figure S6). Generally, using all reads as input allowed both methods to achieve higher recalls but came at a cost to precision. Therefore, henceforth we only show results for Bracken and Kraken2 using all reads, denoted as Bracken(all) and Kraken2(all).

Figure 3 compares TIPP-SD, Kraken2, and Bracken for species detection accuracy using precision-recall curves. Generally, TIPP-SD had better precision and recall than Bracken and Kraken2 with all reads, except for 1000 known species with Illumina reads. For 1000 known species with Illumina reads, Bracken had a broader top right corner, but in many cases, it had lower precision than TIPP-SD for the same recall.

**Figure 3.**
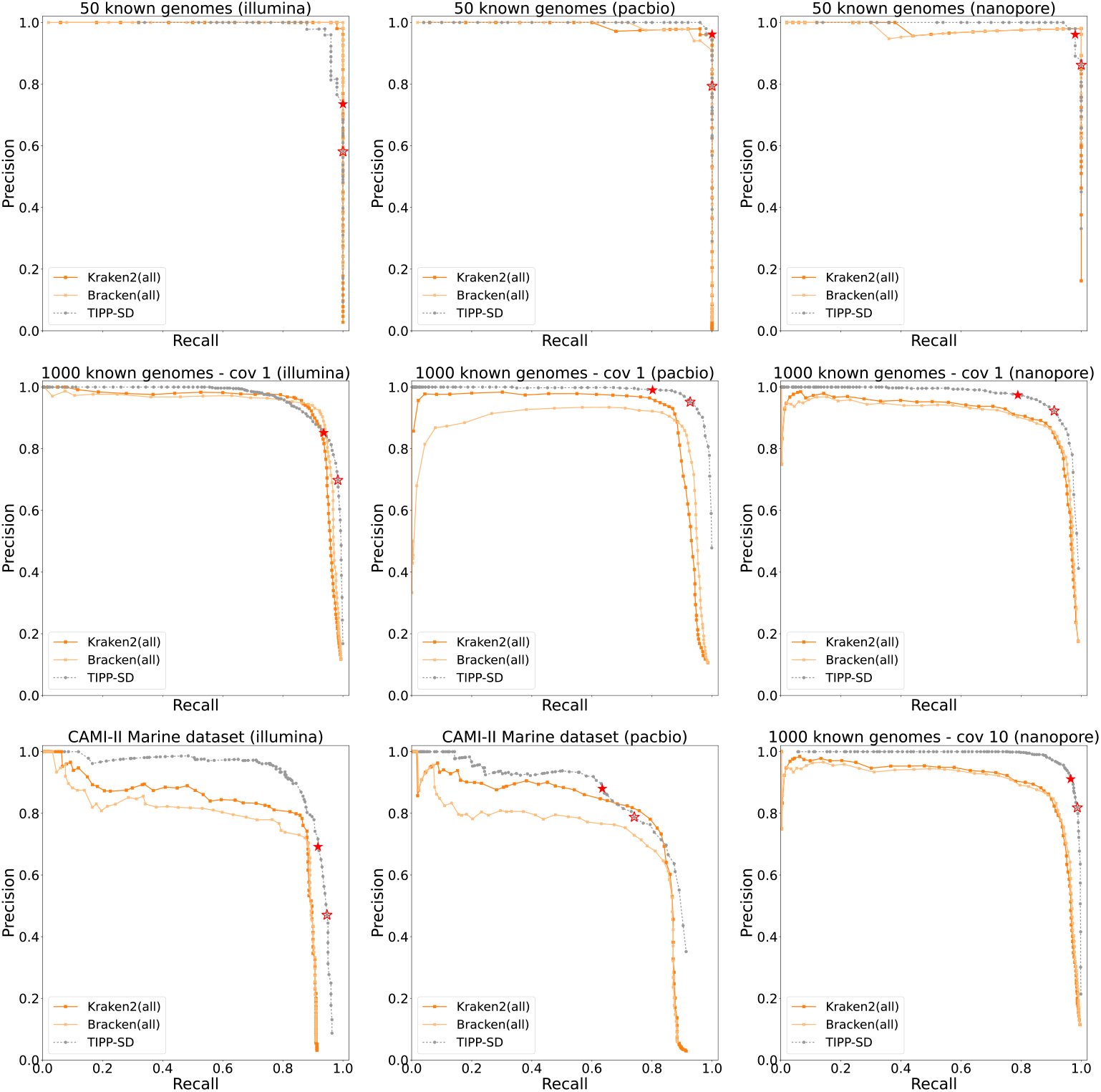
Precision-recall curves of TIPP-SD and Kraken2/Bracken using all reads. The solid and hollow red stars show the precision and recall of TIPP-SD using the “conservative” and “sensitive” thresholds, respectively.

Figures S7 and S8 compare the runtime and memory between TIPP-SD, Bracken, and Kraken2. Kraken2 and Bracken both completed each dataset in under an hour of running time. TIPP-SD was generally slower than Kraken2 and Bracken by about an order of magnitude, but still fast enough to complete on all datasets in 0.5–8.6 hours with 16 CPU cores. TIPP-SD also had a much lower memory usage, using 3–29 GBs compared to ~72 GBs by Kraken2 and Bracken.

### 5.4 Experiment 3: Comparison to Metapresence

Unlike Kraken2, Bracken, and TIPP-SD, Metapresence is based on a small database, containing only 2000 species. To enable a fair comparison (see [6]), we restricted TIPP-SD to a database of the same set of 2000 species as Metapresence, and we refer to the resultant method as TIPP-SD-2000.

We ran Metapresence on the 1000 known species datasets with three types of metagenomic reads (Illumina, PacBio, and Nanopore). We created a Bowtie2 database with genomes of 2000 species, as described earlier in this study.

Figure 4 shows the precision-recall curves of TIPP-SD-2000 and Metapresence on the three 1000 known species datasets. For Illumina reads, TIPP-SD-2000 and Metapresence were similar in accuracy. For PacBio reads, however, Metapresence had lower recall than TIPP-SD-2000, and the highest recall it achieved was around 80%. For Nanopore reads, Metapresence had lower precision for the same recall value when compared to TIPP-SD-2000. At the rightmost side of the precision-recall curve, one can argue that Metapresence achieves a higher precision than TIPP-SD-2000.

**Figure 4.**
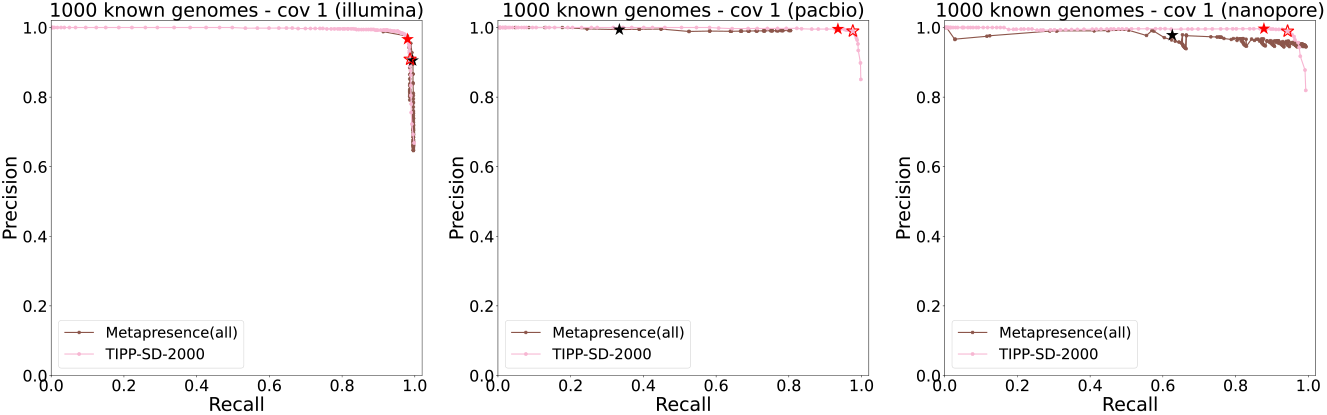
Precision-recall curves of Metapresence and TIPP-SD-2000 (i.e., we use the TIPP3 reference package restricted to the same 2000 species as in Metapresence) on 1000 known species datasets with Illumina, PacBio, and Nanopore reads. The black star in each subfigure shows the precision and recall using the Metapresence default threshold (i.e., a species is considered present if and only if *BER* ≥0.8 and *FUG* ≥0.5). The red solid and hollow stars show the precision and recall of TIPP-SD-2000 using the “conservative” and “sensitive” thresholds, respectively.

Precision and recall of Metapresence using its default threshold values (i.e., *BER* = 0.8, *FUG* = 0.5) are shown in each of the subfigures as black stars. Using this default value, the precision was high for reads of all sequencing technologies, but the recall varied. While recall was very high (close to 100%) for Illumina reads, the recall was low for both PacBio (less than 40%) and Nanopore (less than 65%). Red solid and hollow stars show the precision and recall of TIPP-SD-2000 using the “conservative” and “sensitive” thresholds, respectively. Precision was high for TIPP-SD-2000 while having better recall than Metapresence, except for Illumina reads, where Metapresence achieved a higher recall with lower precision than TIPP-SD-2000’s “conservative” value, and similar performance as TIPP-SD-2000’s “sensitive” value.

We also compared the runtime and memory usage between TIPP-SD-2000 and Metapresence, on the three datasets of 1000 known species with Illumina, PacBio, or Nanopore reads (Figure S9). We observed a noticeable jump in Metapresence’s memory usage when dealing with long reads (e.g., PacBio and Nanopore) compared to short reads (e.g., Illumina). This is likely due to how Bowtie2 handles read alignment against its database, because Bowtie2 is designed for short read alignment. Metapresence is faster than TIPP-SD-2000 on Illumina reads, but slower on both PacBio and Nanopore reads.

### 5.5 Case study: analysis on false-positive species

Here we took a closer look at the detected species by Kraken2, Bracken, and TIPP-SD and tried to understand the properties of their false positives, using the 1000 known species dataset with Illumina, PacBio, and Nanopore reads (center row in Figure 3).

#### False-positive species “closeness” to target species

We looked at the ANI of false-positive species to the “true” species in the 1000 known species dataset with Illumina, PacBio, and Nanopore reads. This study is important to understand if false positives of a method are closely related species to some target species, but have different taxonomic labels.

We used two recall thresholds to extract the list of reported species by each method, at 90% and 95%. We then identified the falsely reported species and computed their ANIs to the closest species among the target species using FastANI. We repeated this process for all tested datasets, but only show the results at 95% for 1000 known species. Results at the 90% recall threshold show similar trends and can be found in the supplementary materials, Figure S10.

At 95% recall (Figure 5) all sequencing technologies show similar trends. TIPP-SD has the smallest number of false positives of all methods, followed by Bracken, and then followed by Kraken2 (which has the largest number). For all sequencing technologies, both Bracken and Kraken2 have false positives across the range of ANI values between 80% and 100% (with Kraken2 having a larger number of false positives at the lower end of this range than Bracken), while TIPP-SD has the majority of its false positives at the upper end of the range. Thus, overall, TIPP-SD shows fewer false positives than the other two methods, and when it has false positives, they tend to be for species that have very similar sequences.

**Figure 5.**
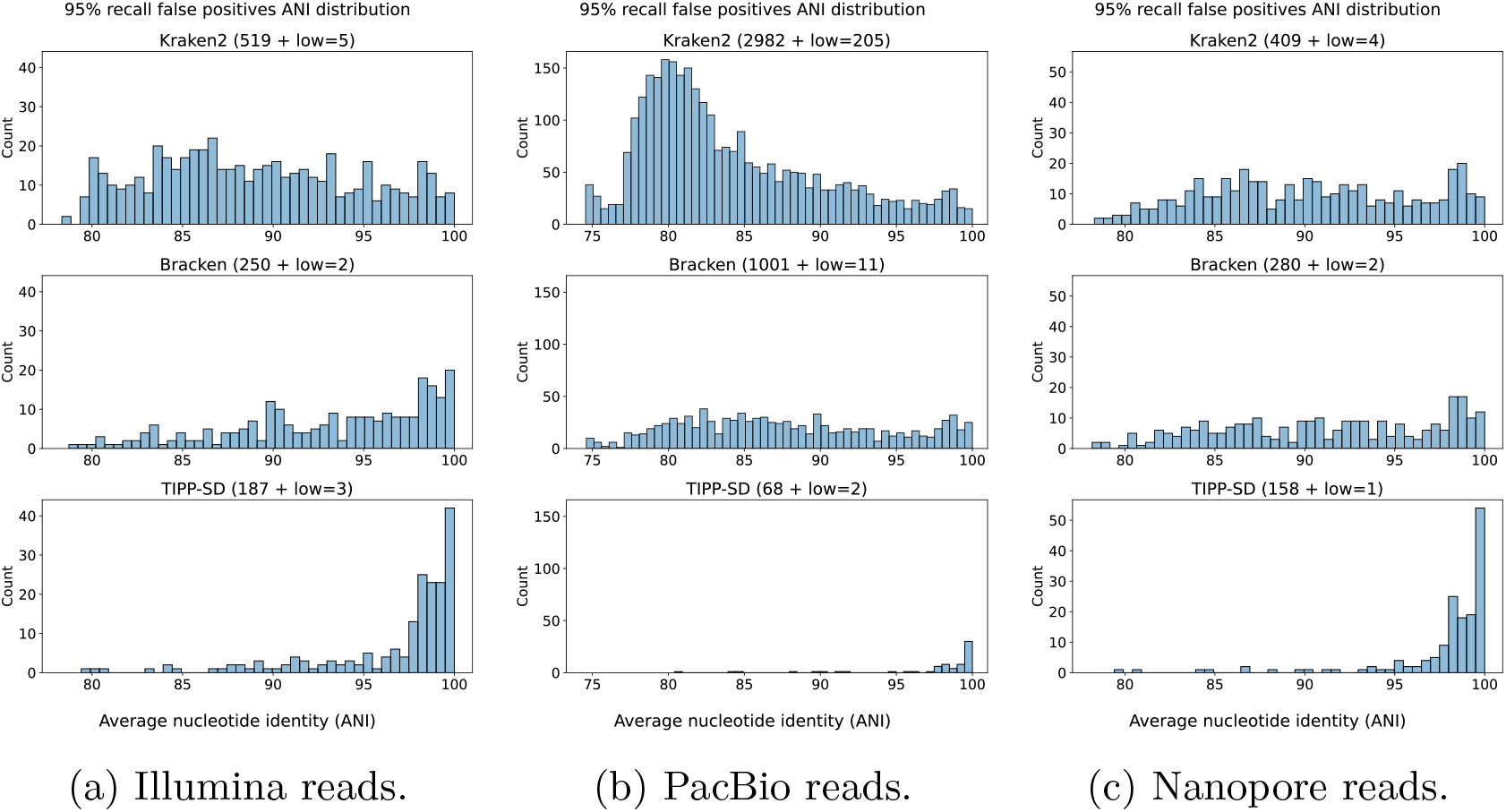
At 95% recall, the distribution of average nucleotide identity (ANI) of false-positive species to their closest species in the set of 1000 known species, for Kraken2, Bracken, and TIPP-SD classifying Illumina (left), PacBio (middle), and Nanopore (right) reads. The total number of false positives for each method is shown in parentheses, with *low* = *X* meaning that there are *X* false positives that do not have any close species in the target (i.e., ≪ 80% ANI according to fastANI).

With this analysis, note that although TIPP-SD may have no obvious advantage in Figure 3 on 1000 known genomes for Illumina reads, this more careful evaluation shows that when TIPP-SD does report false positives, they tend to have high ANI to the closest known species (suggesting that they are very closely related), an advantage that Kraken2 and Bracken do not show here.

#### Initial drop in precision for Bracken and Kraken2

We observed an initial drop in precision for the two methods when analyzing long reads. In the case of PacBio long reads of 1000 known species, this drop in precision is very noticeable for Bracken. We thus examined the top 10 reported species ranked by the number of reads assigned for Bracken and Kraken2 and compared them to the top 10 species reported by TIPP-SD ranked by marker confidence.

Table S2 (supplementary materials) shows the top 10 reported species by Bracken, Kraken2, and TIPP-SD. Kraken2 has only one false positive in set (*Homo sapiens*), Bracken has five false positives (*Homo sapiens* and four others), and TIPP-SD has no false positives. It is noteworthy that both Kraken2 and Bracken reported *Homo sapiens* as one of the detected species, despite that the input reads were only generated from Bacteria and Archaea genomes. Of the four other false positives reported by Bracken, the one with the highest reported threshold is *E. coli*, which has 98.2% ANI to its closest true positive species. All others have lower ANI values to their closest true positive, ranging from 82.5% to 93.0%. In sum, Bracken displayed possibly the most surprising false positives of the three methods.

## 6 Discussion

This study presented TIPP-SD, a new method for species detection. TIPP-SD has high accuracy, with improvements over TIPP3, Kraken2, Bracken, and Metapresence under most tested conditions. The improvement was most noteworthy when there was sequencing error (e.g., PacBio and Nanopore reads) or when some or all species had low abundance (e.g., on the 1000 genomes and on the CAMI-II Marine dataset).

The advantage of TIPP-SD over the competing methods for these two conditions (i.e., reads with sequencing error or low-abundance species) may be due to its ability to use maximum likelihood phylogenetic placement and highly accurate sequence alignments, rather than relying on k-mers. Sequencing depth, or coverage, also has an impact on the methods’ relative and absolute accuracy. Although we only examined Nanopore reads with a coverage of 1 and 10, we observed that TIPP-SD has a better precision-recall curve when the coverage is higher. In addition, its relative accuracy advantage over Bracken and Kraken2 also becomes more noticeable with higher coverage. We also demonstrated that TIPP-SD is fast enough to run on large input data (~36 GBs in FASTA format) in under 10 hours. Overall, TIPP-SD is a competitive method with both reasonable speed and high accuracy for species detection.

The methods also differ in terms of their false positives. At high recall levels, Bracken and Kraken2 often report false-positive species that have low ANIs to any of the target species. TIPP-SD’s false positives generally have very high ANIs (*>* 99%), indicating that they are more likely closely related to some target species but have different taxonomic labels. These “false positives” are hard to eliminate but are also not as undesirable as the ones with low ANIs.

This study suggests several directions for improving TIPP-SD. To improve accuracy in species detection, we could expand the list of “top placements” provided by pplacer to more than its default, which is the top seven. We showed results for TIPP-SD for species detection, but future work should examine results for detecting other taxonomic levels, such as genera, families, or even strains. Although TIPP-SD had very good accuracy, it used the same set of marker genes that were used in TIPP3, and a different set could be considered, given that the purpose of the analysis is different. In particular, we could even consider genes that are not marker genes (i.e., not universal or multi-copy rather than single-copy), given the changed objective. Although TIPP-SD is reasonably fast, a better parallel implementation would speed it up.

Finally, although TIPP-SD did well in this study, its accuracy in comparison to other methods should be explored under additional conditions. While our study showed TIPP-SD had superior accuracy to Kraken2 and Bracken on all datasets where some or all species had low abundance (i.e., 0.1% or less), a more careful analysis of the ability to detect low abundance species is needed. In addition, our evaluation of PacBio reads with high error rates is helpful in understanding robustness to high sequencing error, but the new PacBio reads have much lower sequencing error rates [7, 15]. Similarly, the newer Nanopore technology also has somewhat lower sequencing error than what we examined in our study [8]. We also note that hi-fi reads are increasingly of interest [10]. Thus, future work should examine TIPP-SD under these new sequencing technologies. Other future work includes testing TIPP-SD in comparison to Kraken2 and Bracken on exactly the same set of genomes, and testing TIPP-SD when species in the input sample are not in the TIPP-SD database.

We close with a discussion about the relationship between abundance profiling and species detection, two problems that are very related and both of importance in microbiome analysis. One of the contributions of this study is the observation that techniques that work well for abundance profiling are not necessarily the best for species detection, and vice-versa. For example, [38] established that filtering the input reads to just those that map to selected marker genes (i.e., genes that are expected to be single copy and universal) results in improved estimates of the abundance profile for Kraken2, Bracken, and other methods, but here we have shown that filtering is *detrimental* for Bracken and Kraken2 when used for the species detection problem. Hence, abundance profiling and species detection are different even though related problems, and techniques that work well for one problem may not produce high accuracy for the other problem. Therefore, the development of methods that are specifically designed for each problem is appropriate.

## Acknowledgments

This work was supported in part by NSF grant 2006069 to TW.

## Supplementary Materials

## 1 Additional details on experiment design

### 1.1 TIPP3 reference package raw data

TIPP-SD used the TIPP3 reference package, which was generated from the lists of assemblies (Bacteria and Archaea) downloaded in TIPP2, and marker gene sequences were collected using FetchMG. To download the latest lists of assembly from NCBI, use the following two links:

1. ftp://ftp.ncbi.nlm.nih.gov/genomes/refseq/bacteria/assembly_summary.txt
2. ftp://ftp.ncbi.nlm.nih.gov/genomes/refseq/archaea/assembly_summary.txt

### 1.2 Software commands

All software are given 16 cores and 256 GB of memory to run untill completion.

1 We ran Kraken2 (v2.1.3) with the following command:

~~~
$ kraken2 --db [ Kraken2 database ] \
-- threads 16 -- report [ Kraken2 report ] \
[ query reads file ] > [ Kraken2 output ]
~~~

2 We ran Bracken (v2.9) with the following command (using Kraken2 output):

~~~
$ bracken -d [ Bracken database ] -i [ Kraken2 report ] \
-w [ Bracken report ] -t 10 -r 150 \
-o [ Bracken output ]
~~~

3 We ran TIPP-SD and its variants with the following commands. --`alignment-method` controls what method we used to align the reads, and --`placement-method` and --`bscampp-mode` control what method we used for read placement.

~~~
$ run_tipp3.py abundance -i [ query reads file ] \
-r [ reference package directory ] -- outdir [ output directory ] \
--alignment - method XXX \
--placement - method YYY \
--bscampp - mode ZZZ \
-t 16
~~~

4 For Metapresence (v1.0), we first aligned the input reads using Bowtie2 (v2.5.4) and samtools (v1.19.2):

~~~
$ bowtie2 -f [ query reads file ] -x [ bowtie2 database index ] -p 16 \
| samtools view -b -@ 16 \
| samtools sort -@ 16 > [bam alignment file ]
~~~

We then indexed the bowtie2 alignment with samtools:

~~~
$ samtools index -@ 16 [bam alignment file ]
~~~

Finally, we ran metapresence to analyze the alignment results:

~~~
$ metapresence.py [ reference genomes directory ] [ bam alignment file ] \
~~~

-o [ output prefix ] -p 16

5 To compute average nucleotide identity (ANI) between two sets of genomes, we used fastANI (v1.34) and the following command. Note that fastANI does not return comparisons that have ≪ 80% ANI.

~~~
$ fastANI --ql [ query list of genomes ] --rl [ target list of genomes ] \
-o [ output file ] -t 16
~~~

**Table S1:**
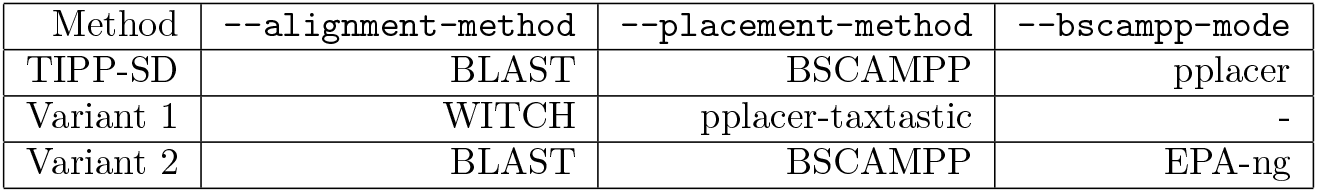
Parameter settings for each TIPP-SD variant.

## 2 Additional results

### 2.1 Additional results for Experiment 1

#### 2.1.1 TIPP-SD vs. TIPP3

**Figure S1:**
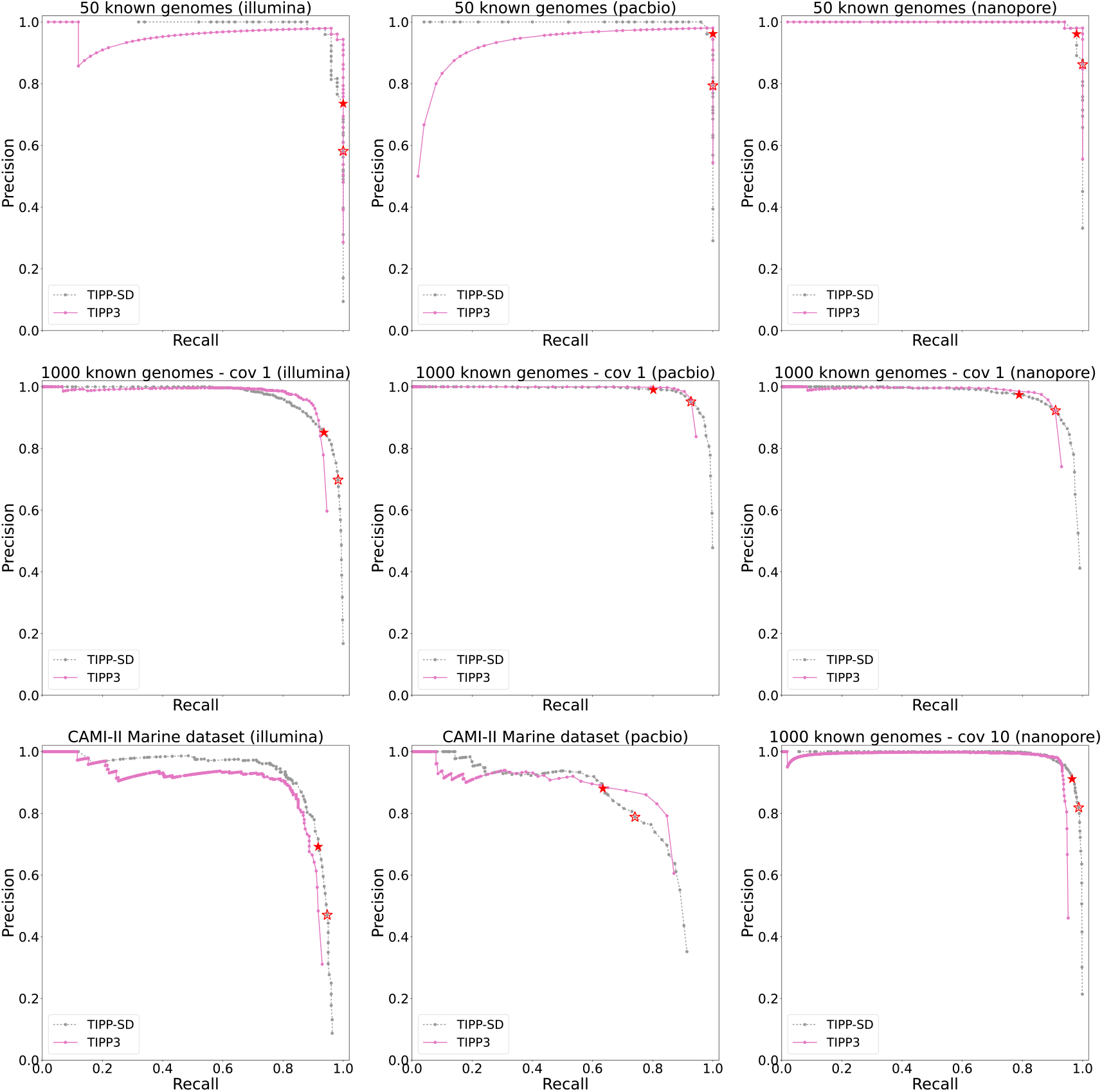
Precision/recall of TIPP-SD compared to TIPP3. TIPP3’s precision/recall curve is obtained by setting thresholds on the abundance level, and only species with an abundance greater than the threshold are considered as “detected” by TIPP3. Red stars mark the precision and recall values of using the “conservative” (solid) and “sensitive” (hollow) threshold values for TIPP-SD. While TIPP3 matches the precision and recall of TIPP-SD for the 50 genome case, it is less accurate for larger numbers of genomes.

#### 2.1.2 TIPP-SD variants

**Figure S2:**
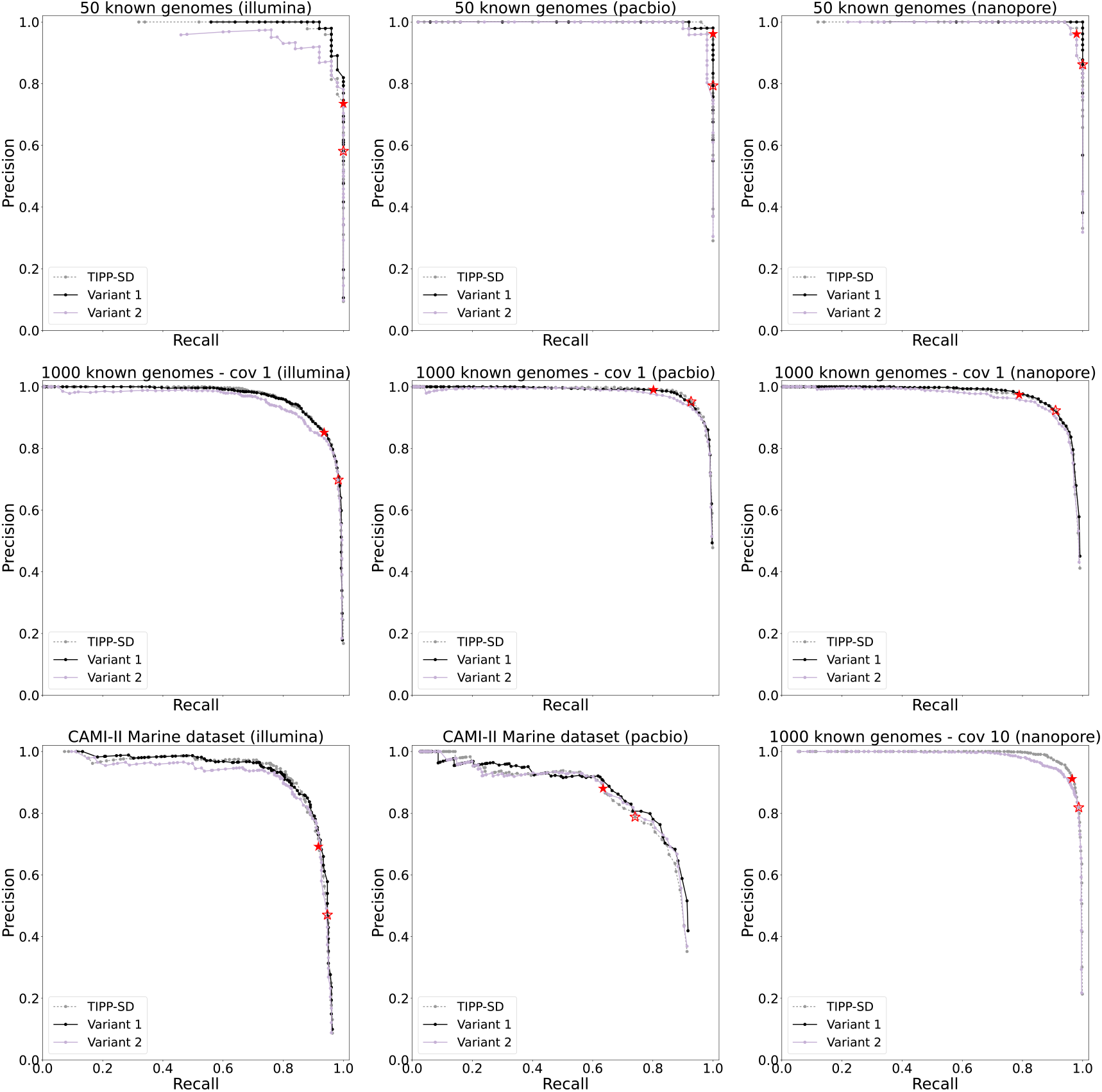
Precision/recall of TIPP-SD, Variant 1, and Variant 2 using marker confidence as the species detection strategy. Variant 1 was not run for Nanopore reads of 1000 known species with a coverage of 10, due to the long running time. Red stars mark the precision and recall values of using the “conservative” (solid) and “sensitive” (hollow) threshold values for TIPP-SD. See Table S1 for definitions of TIPP-SD variants.

**Figure S3:**
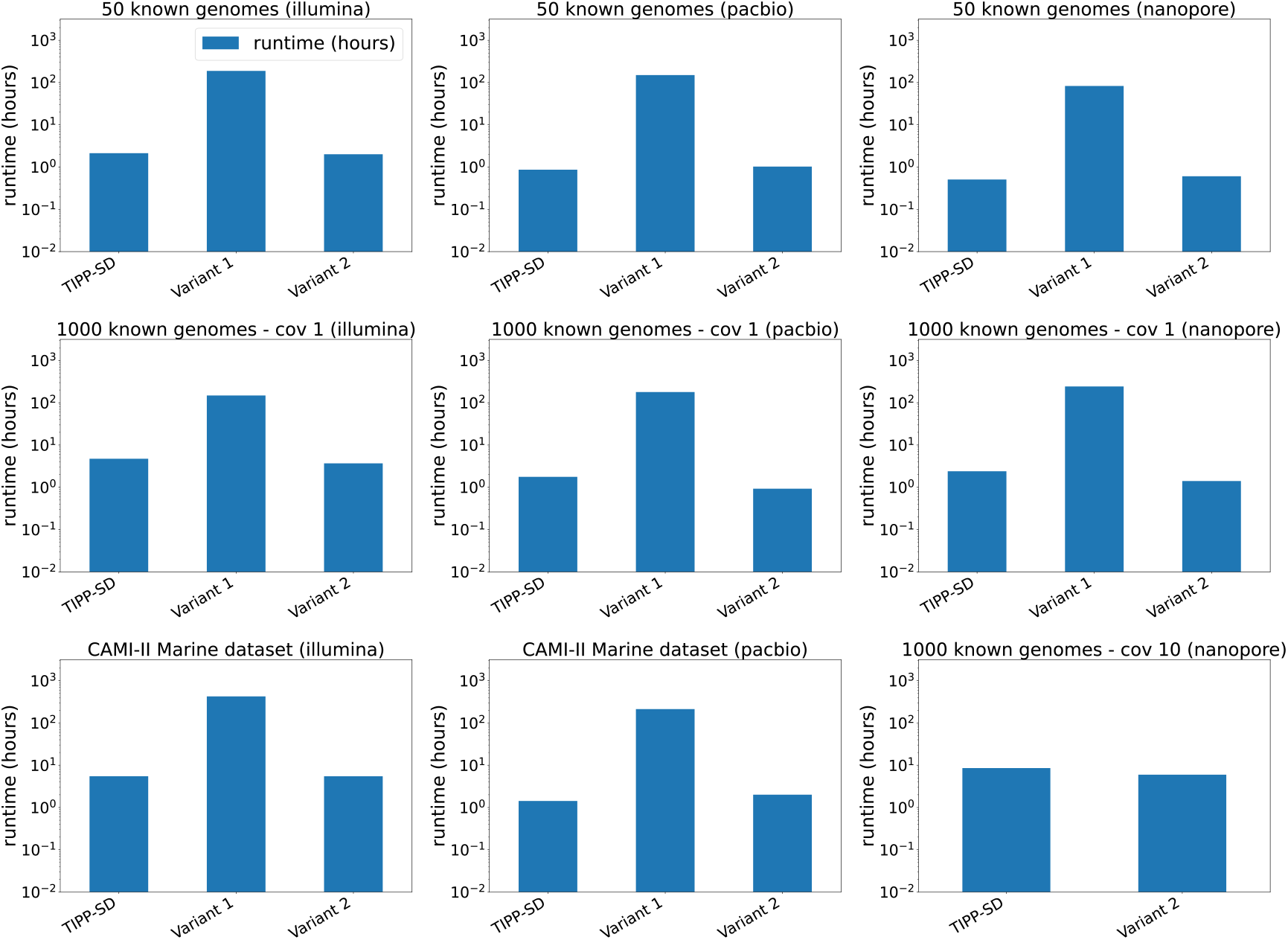
Runtime in hours (in log-scale) of TIPP-SD, Variant 1, and Variant 2. Variant 1 was not run for Nanopore reads of 1000 known species with a coverage of 10, due to the long running time.

**Figure S4:**
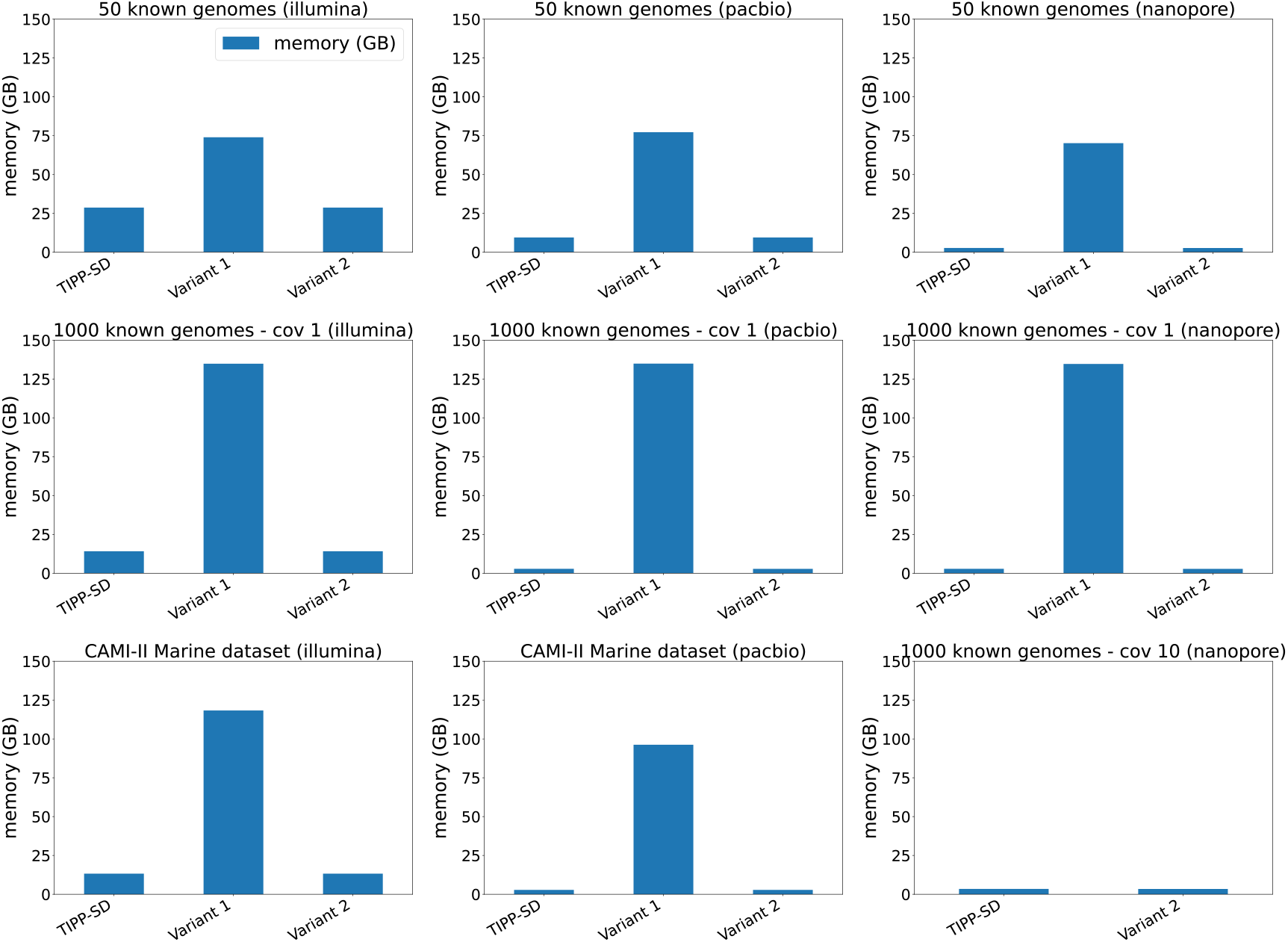
Memory usage in GBs of TIPP-SD, Variant 1, and Variant 2. Variant 1 was not run for Nanopore reads of 1000 known species with a coverage of 10, due to the long running time.

#### 2.1.3 Selecting default detection threshold

Figure S5 shows the marker confidence to F1 score of TIPP-SD, Variant 1, and Variant 2 on the tested datasets. For the easy datasets (i.e., high coverage and/or Illumina reads), a wide range of marker confidence values leads to high F1 scores.

Generally, for 50 known genomes, the peak F1 score happened when marker confidence was 0.3– 0.6 for Illumina reads, 0.2–0.3 for PacBio reads, and 0.2–0.3 for Nanopore reads. For 1000 known genomes with a coverage of 1, the peak F1 score happened when marker confidence was 0.2–0.3 for Illumina, 0.1–0.2 for PacBio, and 0.1–0.2 for Nanopore reads. For 1000 known genomes with Nanopore reads and a coverage of 10, the range of marker confidence for peak F1 score widened to 0.2–0.4. Finally, for CAMI-II Illumina reads, the marker confidence range is 0.2–0.5, and for PacBio reads, the range is 0.1–0.2.

The overall trends suggest that a threshold around 0.2 should be accurate for most conditions. When the dataset is “easy” (e.g., Illumina reads or high coverage), larger thresholds (e.g., around 0.6) are more suitable, but when the dataset is “difficult” (e.g., low coverage or Pacbio reads), then a lower threshold (e.g., around 0.2) is needed.

**Figure S5:**
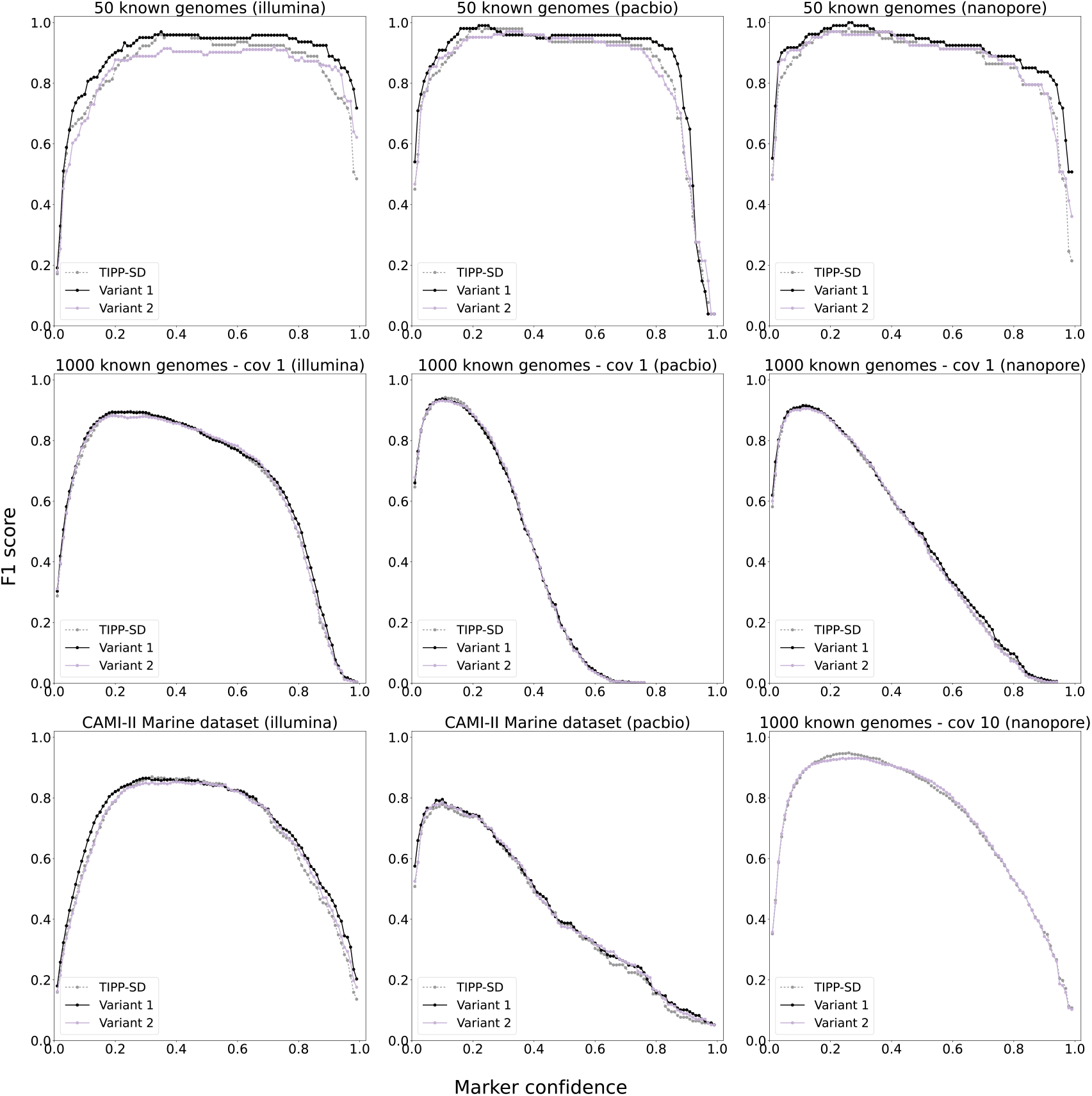
Marker confidence (x-axis) to F1 score (y-axis) of TIPP-SD, Variant 1, Variant 2 on the examined datasets. F1 score is computed as 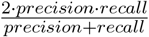, a measurement between 0 and 1 on the quality of both precision and recall. Variant 1 was not run for Nanopore reads of 1000 known species with a coverage of 10, due to the long running time.

### 2.2 Additional results for Experiment 2

#### 2.2.1 Filtering vs. all reads

Figure S6 compares Kraken2 and Bracken when using all or filtered reads as input for species detection. When dealing with 50 known genomes, both Kraken2 and Bracken with filtered reads have very high accuracy, showing almost perfect precision/recall curves. Kraken2 and Bracken, with all reads on 50 known genomes, have a drop in precision for the same level of recall when compared to their variants with filtered reads.

However, the results with 1000 species with a coverage of 1 are slightly different. Generally, both Kraken and Bracken achieve better recall with all reads as input than with filtered reads. Precision is slightly higher for using filtered reads at low levels of recall, but drops at higher values. Comparing a coverage of 1 and 10 for the Nanopore reads, Kraken2(filtered) and Bracken(filtered) improve in their recall, but the impact on the performance of Kraken2(all) and Bracken(all) is not as noticeable.

For CAMI-II datasets, for Illumina reads, Kraken2(filtered) improves upon Kraken2(all) in terms of precision, but Bracken(filtered) and Bracken(all) have similar performance for both precision and recall. For PacBio reads, Bracken(filtered) has high precision but low recall. Kraken2(all) achieves higher precision than Bracken(all) with similar recall.

In conclusion, for the task of species detection, if the focus is to achieve a high recall, we recommend running with all reads as inputs for Bracken and Kraken2, as it generally leads to better recall.

**Figure S6:**
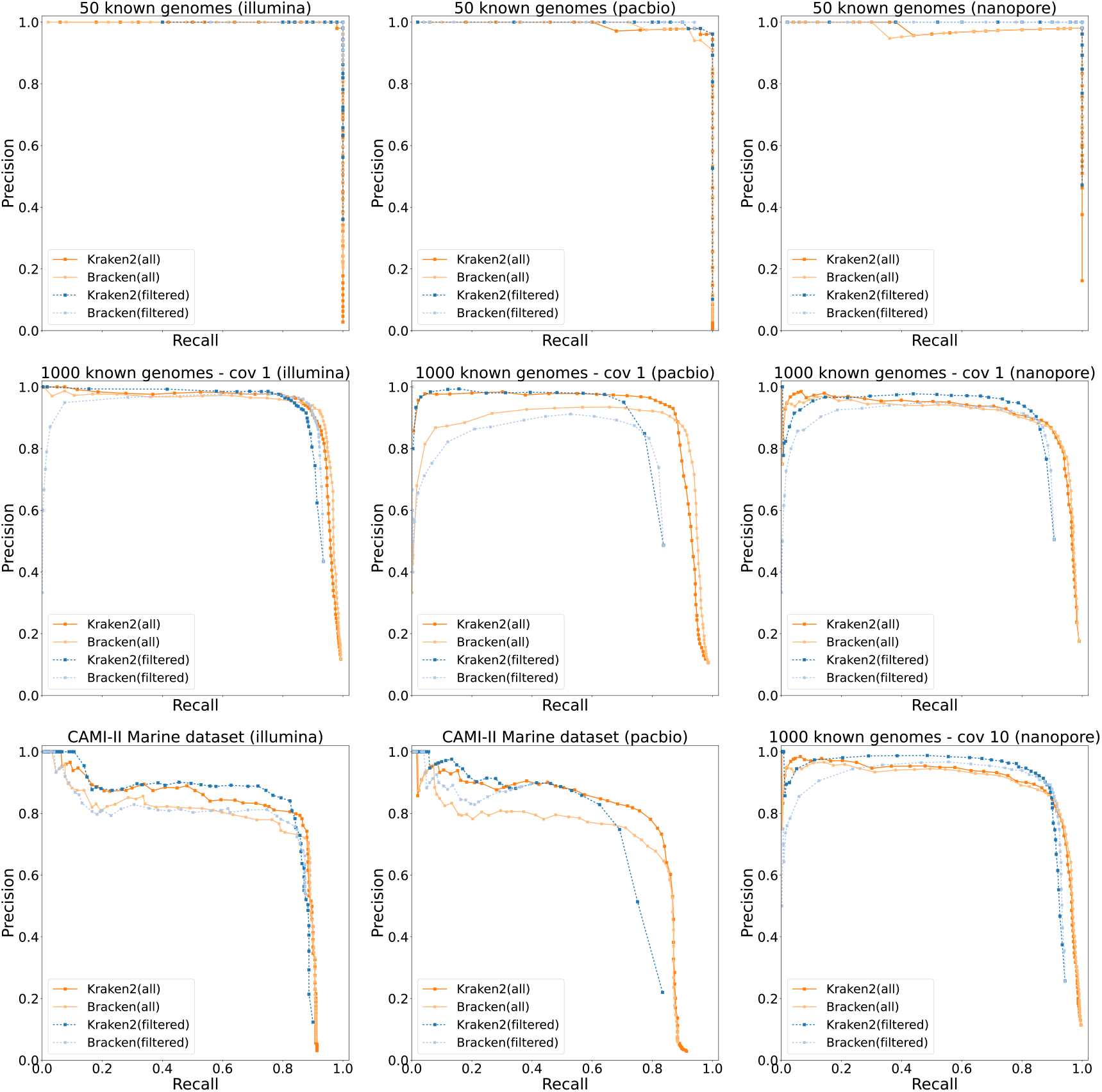
Precision/recall of Kraken2 and Bracken using all or filtered reads as input. For filtered reads, the lines are dashed.

#### 2.2.2 Runtime and memory

**Figure S7:**
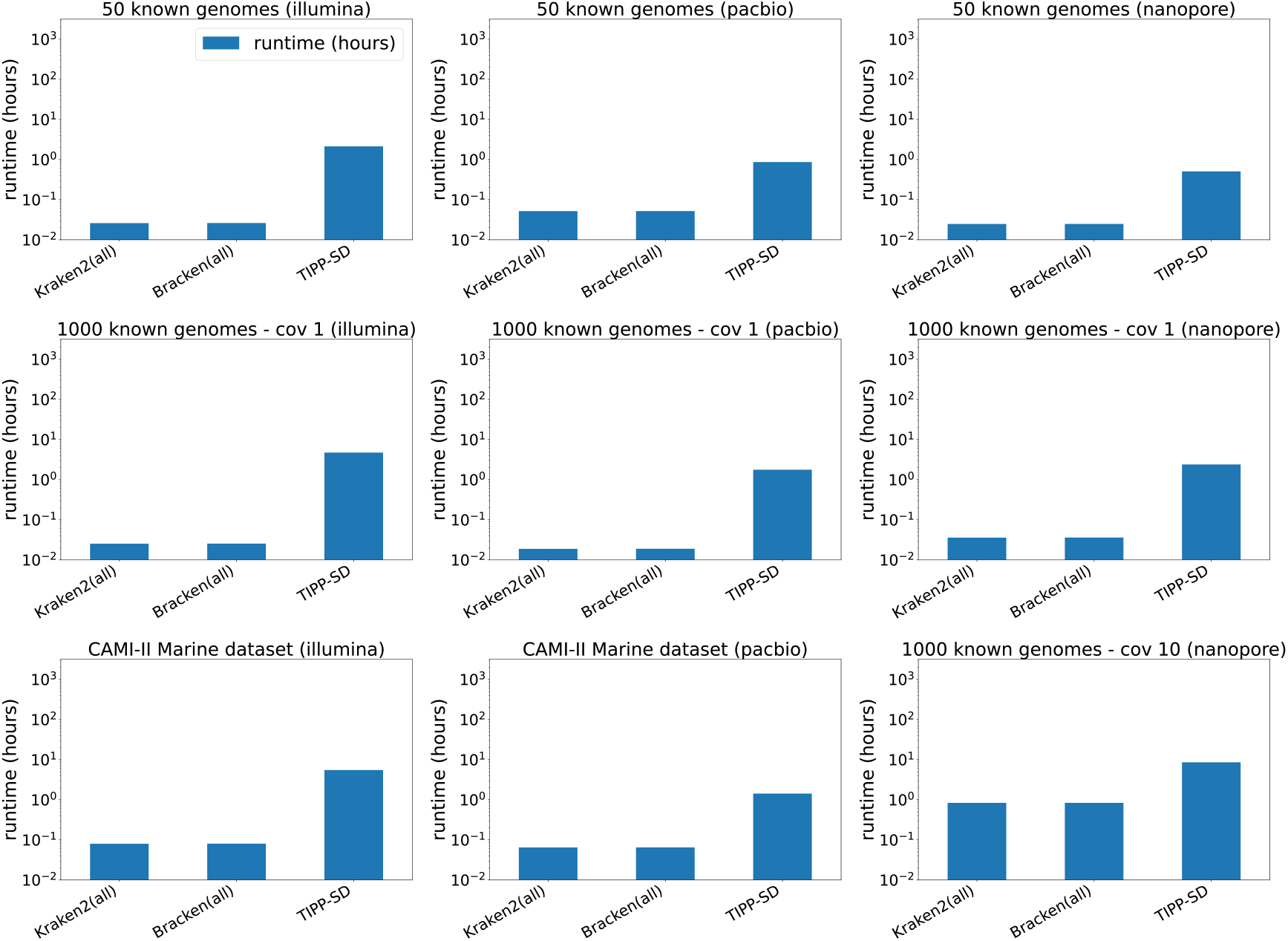
Runtime in hours (in log-scale) of TIPP-SD and Kraken2/Bracken using all reads.

**Table S2:** The top 10 reported species of Kraken2, Bracken, and TIPP-SD when classifying PacBio reads of 1000 known species. The “reported threshold” for Kraken2 and Bracken is ranked by the number of reads assigned to a species, and for TIPP-SD, this is the marker confidence value. The “presence” column denotes whether the species is a true positive.

| Kraken2 |  |  |  | Bracken |  |  |  | TIPP-SD |  |  |  |
| --- | --- | --- | --- | --- | --- | --- | --- | --- | --- | --- | --- |
| Species | Taxid | Reported Threshold | Presence | Species | Taxid | Reported Threshold | Presence | Species | Taxid | Reported Threshold | Presence |
| H. sapiens | 9606 | 6973 | FALSE | P. larvae | 1464 | 28749 | TRUE | D. nodosus | 870 | 0.74 | TRUE |
| S. acidiphila | 466153 | 1922 | TRUE | E. coli | 562 | 10301 | FALSE | L. pinisoli | 2589080 | 0.71 | TRUE |
| Streptomyces sp. YIM 121038 | 2136401 | 1854 | TRUE | H. sapiens | 9606 | 8462 | FALSE | Verrucomicrobium sp. GAS474 | 1882831 | 0.67 | TRUE |
| S. griseochromogenes | 68214 | 1838 | TRUE | D. acidovorans | 80866 | 5130 | TRUE | V. quintilis | 1117707 | 0.66 | TRUE |
| M. ossetica | 1882682 | 1810 | TRUE | B. ambifaria | 152480 | 3264 | FALSE | P. borealis | 1387353 | 0.66 | TRUE |
| T. plasticadhaerens | 2527974 | 1809 | TRUE | Rhizobium sp. N1341 | 1703962 | 3251 | TRUE | M. bathoardescens | 1301915 | 0.66 | TRUE |
| N. koreensis | 354356 | 1794 | TRUE | P. aeruginosa | 287 | 2911 | FALSE | P. inopinatus | 114090 | 0.65 | TRUE |
| S. venezuelae | 54571 | 1721 | TRUE | B. cereus | 1396 | 2523 | TRUE | N. ponticola | 2496866 | 0.65 | TRUE |
| S. espanaensis | 103731 | 1690 | TRUE | Streptomyces sp. CRLD-Y-1 | 2964608 | 2455 | FALSE | C. papalotli | 1208583 | 0.65 | TRUE |
| Actinoplanes sp. N902-109 | 649831 | 1661 | TRUE | S. venezuelae | 54571 | 2146 | TRUE | Pigmentiphaga aceris | 1940612 | 0.63 | TRUE |

**Figure S8:**
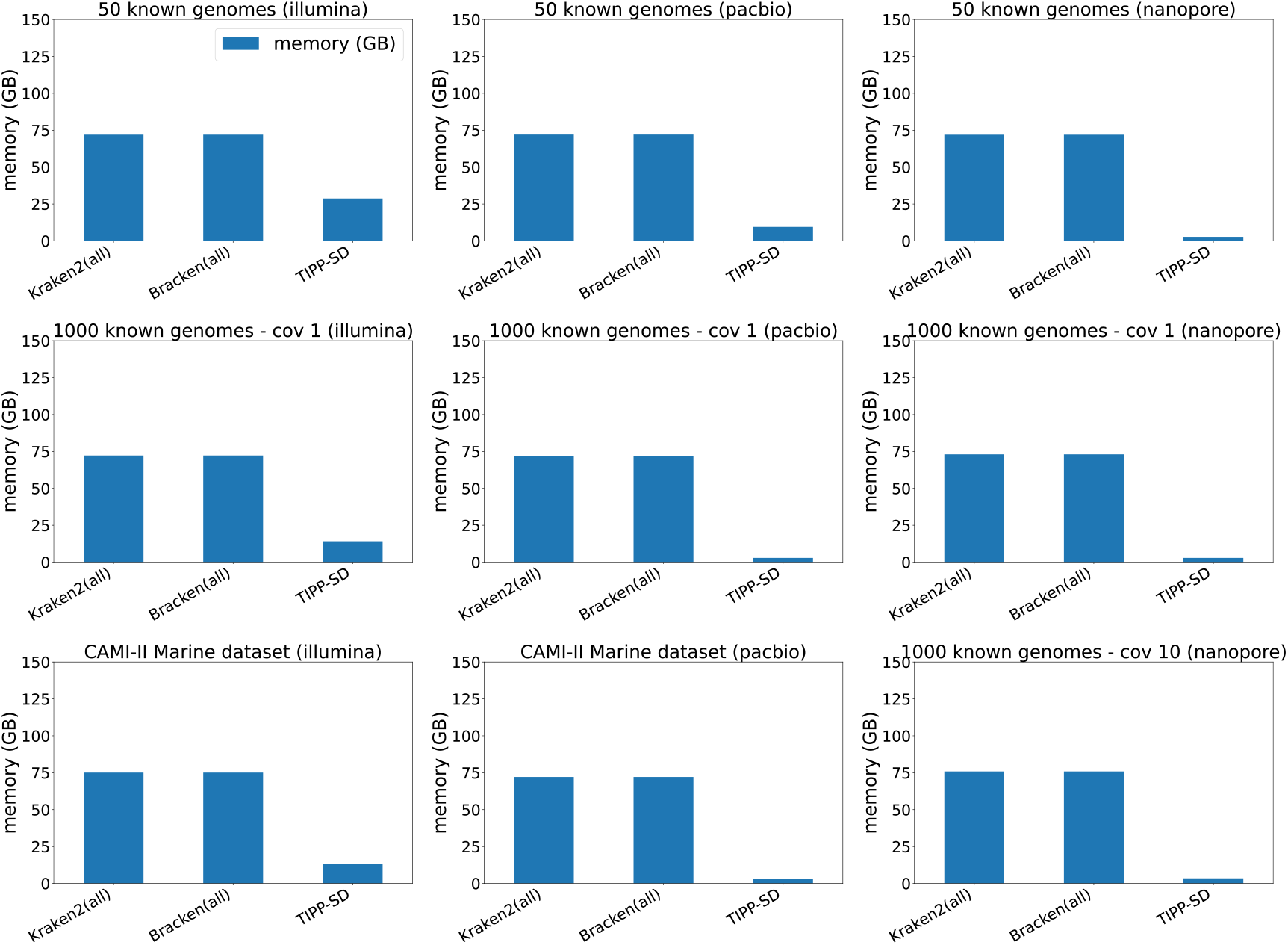
Memory usage in GBs of TIPP-SD and Kraken2/Bracken using all reads.

### 2.3 Additional results for Experiment 3

#### 2.3.1 Runtime and memory

**Figure S9:**
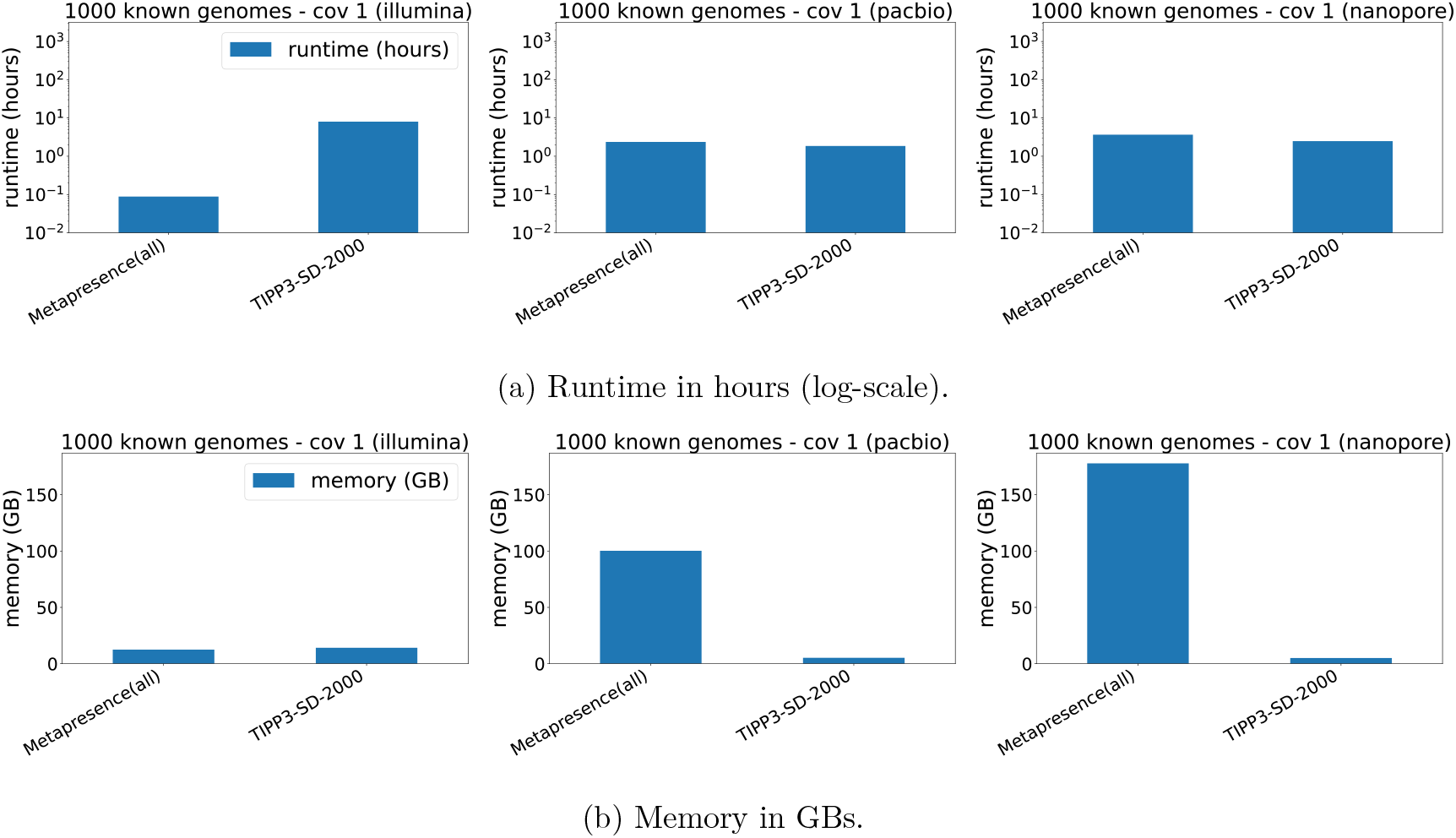
Runtime and memory usage of TIPP-SD-2000 and Metapresence on 1000 known species datasets with Illumina, PacBio, and Nanopore reads. Coverages for the 1000 species datasets are 1.

### 2.4 Additional results for the case study

**Figure S10:**
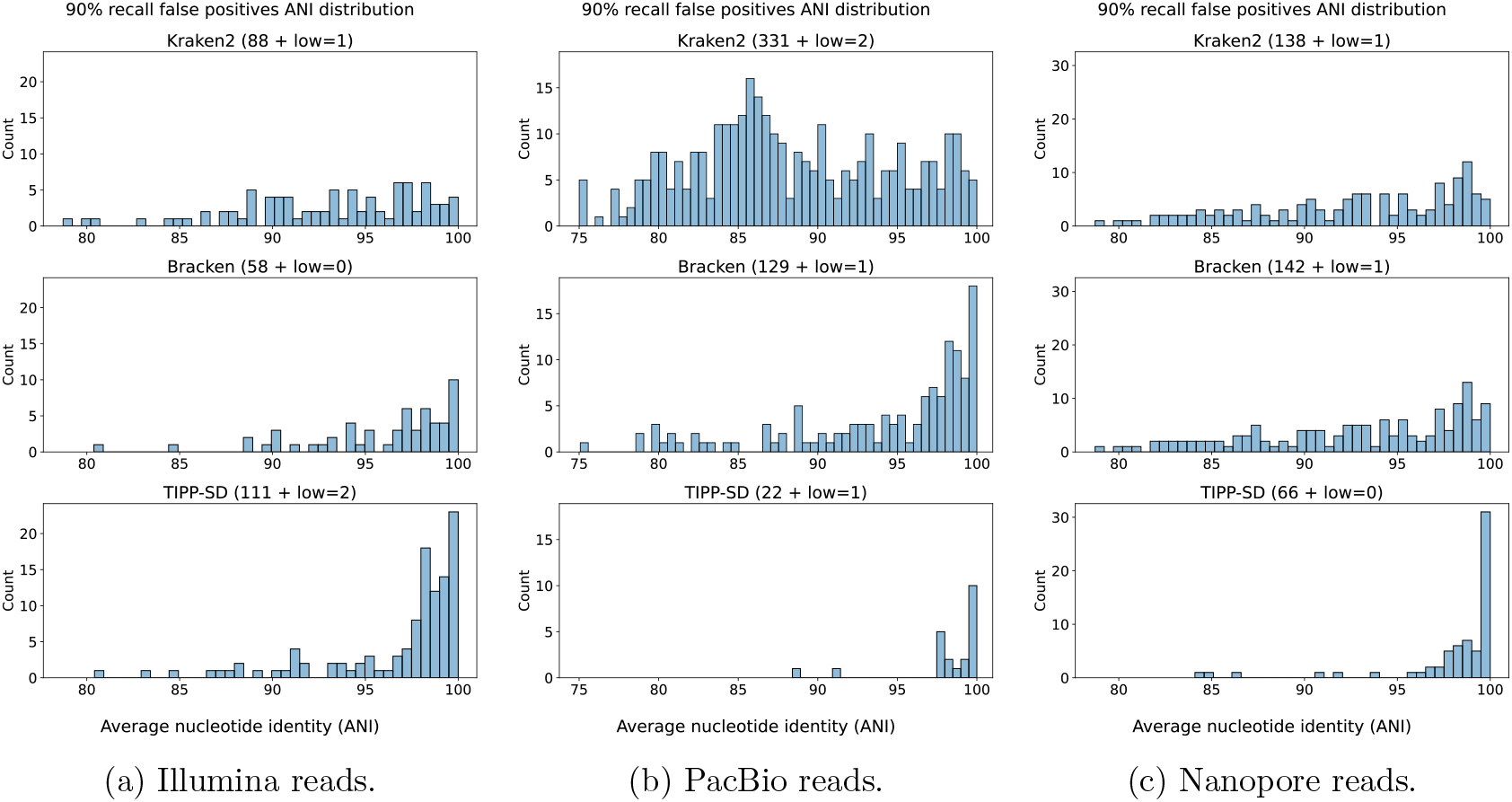
At 90% recall, the distribution of average nucleotide identity (ANI) of false-positive species to their closest species in the set of 1000 known species, for Kraken2, Bracken, and TIPP-SD classifying Illumina (left), PacBio (middle), and Nanopore (right) reads. The total number of false positives for each method is shown in parentheses, with *low* = *X* meaning that there are *X* false positives that do not have any close species in the target (i.e., ≪ 80% ANI according to fastANI).

